# Linking patterns of genetic variation to processes of diversification in Malaysian torrent frogs (Anura: Ranidae: *Amolops*): a landscape genomics approach

**DOI:** 10.1101/628891

**Authors:** Kin Onn Chan, Rafe M. Brown

**Affiliations:** Biodiversity Institute and Department of Ecology and Evolutionary Biology, University of Kansas, Lawrence, Kansas, 66045, USA; Lee Kong Chian Natural History Museum, Faculty of Science, National University of Singapore, 2 Conservatory Drive, Singapore 117377

**Keywords:** isolation-by-colonization, isolation-by-distance, isolation-by-environment, redundancy analysis, landscape genetics, population genetics, dbRDA

## Abstract

The interplay between environmental attributes and evolutionary processes can provide valuable insights into how biodiversity is generated, partitioned, and distributed. This study investigates the role of spatial, environmental, and historical factors that could potentially drive diversification and shape genetic variation in Malaysian torrent frogs. Torrent frogs are ecologically conserved, and we hypothesize that this could impose tight constraints on dispersal routes, gene flow, and consequently genetic structure. Moreover, levels of gene flow were shown to vary among populations from separate mountain ranges, indicating that genetic differentiation could be influenced by landscape features. Using genome-wide SNPs in conjunction with landscape variables derived from GIS, we performed distance-based redundancy analyses and variance partitioning to disentangle the effects of isolation-by-distance (IBD), isolation-by-environment (IBE), and isolation-by-colonization (IBC). Our results demonstrated that IBE, contributed minimally to genetic variation. Intraspecific population structure can be largely attributed to IBD, whereas interspecific diversification was primarily driven by IBC. We also detected two distinct population bottlenecks, indicating that speciation events were likely driven by vicariance or founder events.

## INTRODUCTION

Genetic variation forms the building blocks on which evolution acts on and plays a crucial role in the diversification and adaptability of a species (Dobzhansky, 1937; Lewontin, 1974; Mayr, 1954). An important evolutionary process that underlies genetic variation among populations is gene flow. Barriers to gene flow can be influenced by intrinsic factors such as reproductive incompatibility (Seehausen *et al*., 2014), extrinsic factors such as physical barriers or ecological constraints (Edwards, Keogh, & Knowles, 2012; Geissler *et al*., 2015), or a combination of both whereby local adaptation to disparate environments produces differential reproductive success (Glor & Warren, 2011; Nosil, 2012; Savolainen, Lascoux, & Merilä, 2013). Understanding the factors that affect gene flow among populations can therefore provide valuable insights into how biodiversity is generated, maintained, partitioned, and distributed—and these may be critically important for addressing classic evolutionary questions related to speciation (Chan *et al*., 2017; Jackson *et al*., 2017), adaptation (Nadeau *et al*., 2016), hybridization (Payseur & Rieseberg, 2016), and/or conservation (Cushman *et al*., 2006; Castillo *et al*., 2014).

The field of landscape genetics focuses on the influence of landscape characteristics on gene flow and consequently, genetic differentiation across the geographical template (Sork *et al*., 1999; Manel *et al*., 2003; Storfer *et al*., 2007). Populations can diverge through selection or local adaptation to different environments, a process generally referred to as isolation-by-environment, IBE (Wang & Summers, 2010; Wang & Bradburd, 2014). Isolation-by-environment can be caused by abiotic factors such as climatic variables (Leamy *et al*., 2016), or biotic factors such as vegetation density and host (Via & Hawthorne, 2002). However, genetic differentiation can also be shaped by neutral processes such as random genetic drift, where the rate of gene exchange in continuous populations inhabiting a homogenous habitat is dependent upon the distance among populations. Under this scenario, populations that are geographically close to each other tend to be more genetically similar than populations that are far apart, a phenomenon known as isolation-by-distance, IBD (Wright, 1943). Isolation-by-resistance, IBR (Mcrae, 2006) on the other hand, produces genetic differentiation as a result of landscape features such as habitat corridors, mountains, and rivers that form physical barriers or resistances to migration (Spear *et al*., 2010; Cushman & Landguth, 2012; Ozerov *et al*., 2012; Cushman, Lewis, & Landguth, 2014). Additionally, historical events such as vicariance or founder events/colonization (isolation-by-colonization, IBC) can also generate genetic structure that persists through time, independent of landscape/environmental characteristics (Nason, Hamrick, & Fleming, 2002; Orsini *et al*., 2013b; Dewoody, Trewin, & Taylor, 2015; Lanier *et al*., 2015; Nadeau *et al*., 2016). Assessing the relative contribution of historical, environment and geographic factors in shaping genetic variation is a key focus of landscape genetics—with a number of studies suggesting that environmental adaptation may play an important role in population divergence (Sexton, Hangartner, & Hoffmann, 2014; Leamy *et al*., 2016).

Disentangling adaptive (IBE) from neutral processes (IBD, IBC) can be challenging (Wang & Bradburd, 2014). For example, patterns of genetic differentiation generated by IBD can be similar to that of IBE when geography is correlated with environmental variation (Meirmans, 2012). Distinguishing patterns generated by founder events (IBC) from IBD and IBE can also be complicated because they can be influenced by several processes: local adaptation can reinforce founder events resulting in patterns similar to IBE (De Meester *et al*., 2002), and colonization can produce allele frequency gradients similar to IBD or IBE because colonization routes often covary with environmental gradients (Orsini *et al*., 2013a; Nadeau *et al*., 2016). Because selective environmental gradients, geography, and colonization routes are often spatially correlated, decoupling the relative effects of adaptive from neutral processes can be problematic.

Peninsular Malaysia is one of the most biodiverse yet highly threatened regions in Southeast Asia (Myers *et al*., 2000). New species are being discovered at an unprecedented rate (Chan *et al*., 2009; Chan, Grismer, & Grismer, 2011; Chan *et al*., 2014, 2018a; Chan & Ahmad, 2010; Grismer *et al*., 2012, 2014b, 2017; Quah *et al*., 2017) and many more await discovery (Chan, Grismer, & Brown, 2018b; Chan & Grismer, 2019). Unfortunately, deforestation and habitat loss (Sodhi *et al*., 2004; Hansen *et al*., 2013) are occurring at a similarly-increasing pace. Despite this imminent crisis, no studies of vertebrates have yet investigated processes involved in shaping genetic diversity across the region’s highly heterogeneous landscape. Recent phylogenetic studies have shown that numerous species of amphibians and reptiles are endemic to specific mountain ranges (Wood *et al*., 2009; Grismer *et al*., 2012, 2013, 2015, Chan *et al*., 2014, 2018a; Sumarli *et al*., 2016), indicating that landscape characteristics may play an important role in shaping genetic diversity. In this study, we analyze genome-wide SNP data within a landscape genomic context, to investigate the factors that shape genetic differentiation in Malaysian torrent frogs (genus *Amolops*).

Torrent frogs are widely distributed across Asia from northeastern India, southern China, and southwards throughout mainland Southeast Asia (Frost, 2018). Throughout their range, torrent frogs are highly specialized and ecologically conserved – only occurring in torrential zones of boulder-strewn forest streams. Adults and tadpoles possess adaptive morphological characters (expanded toe pads in adults; oral disc and abdominal sucker in tadpoles), and larval development is dependent on the high concentration of dissolved oxygen provided by fast-moving water (Nodzenski & Inger, 1990; Pham *et al*., 2015). We hypothesize that these strict ecological requirements could potentially impose tight constraints on dispersal routes, gene flow, and consequently genetic structure, especially when contextualized across a highly heterogeneous geographic template such as Peninsular Malaysia.

The landscape of Peninsular Malaysia (PM) is largely characterized by mountain ranges that transect the peninsula longitudinally. Three species of torrent frogs widely occur along river networks throughout these mountain ranges, but not in the intervening flatlands where habitat is unsuitable. *Amolops larutensis* occurs on the Bintang and Titiwangsa ranges along the western half of PM (shaded orange in Fig. 1), while *A. gerutu* and *A. australis* occur on the Eastern (shaded purple) and Southern ranges (shaded blue) respectively along the eastern region of PM (Fig. 1; Chan, Abraham, et al., 2018). Gene flow was shown to occur among populations of *A. larutensis* from the Bintang and Titiwangsa ranges but *A. gerutu* and *A. australis* were genetically isolated (Chan *et al*., 2017). Additionally, *A. larutensis* and *A. gerutu* were found in sympatry without interbreeding at one locality where the Titiwangsa and Eastern ranges contact (Fig. 1). This alludes to a complex scenario of divergence and secondary contact that may be influenced (in part) by landscape variables, especially mountain ranges. Therefore, the objectives of this study are to: (i) test the prediction that the spatial configuration of mountain ranges may play a role in partitioning genetic variation (Mountain Range Hypothesis); (ii) identify the relative contributions of other factors that may shape genetic diversity, including geographic distance (IBD), river basins, forest cover, habitat suitability (IBE), and historical vicariance/founder events (IBC); and (iii) determine the underlying evolutionary processes that are associated with the observed patterns of genetic differentiation. This study demonstrates how landscape and population genomic methods can be used in concert to elucidate the underlying mechanisms that generate, maintain, and distribute biodiversity.

**Fig. 1.**
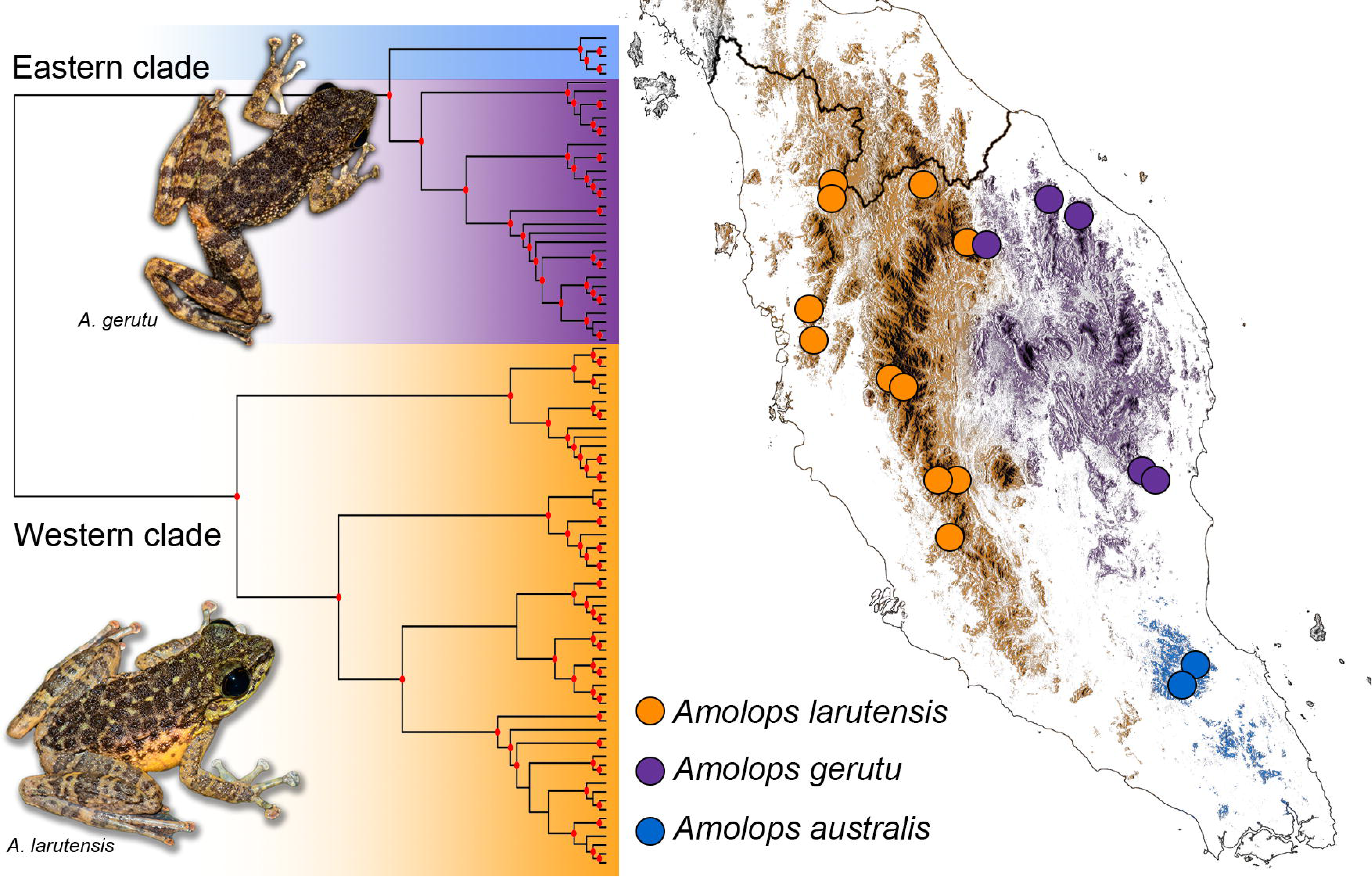
Distribution of genetic samples across the mountain ranges in Peninsular Malaysia and a maximum likelihood (ML) cladogram based on 17,123 SNPs adapted from Chan *et al*., (2017). Red circles on the phylogeny represent ML bootstrap support values > 95. Mountain ranges are highlighted to match the major clades: orange=Bintang+Titiwangsa range; purple=eastern range; blue=southern range.

## MATERIAL AND METHODS

### Genetic sampling and landscape genomic data

We used the genomic dataset generated by Chan *et al*., (2017), which comprised 95 individuals from all three species of *Amolops* that occur in Peninsular Malaysia (*A. larutensis, A. gerutu*, and *A. australis;* Chan et al., 2018). Samples were obtained from 20 unique localities across all major mountain ranges in Peninsular Malaysia and genotyped at 17,123 unlinked SNP loci (Fig. 1). Details on genomic sequencing, data assembly, and species delimitation are beyond the scope of this study and can be obtained from Chan et al. (2017, 2018).

Genetic distances were represented by pairwise population F_st_ values that were calculated using GENODIVE v2.0b27 (Meirmans & Van Tienderen, 2004). Due to the high level of interspecific divergence, missing data were imputed using population allele frequencies prior to F_st_ calculations. Geographic distances were transformed into distance-based Moran’s eigenvector maps (dbMEM) for use as an independent spatial variable in subsequent ordination analyses (Legendre, Fortin, & Borcard, 2015). The dbMEM analyses was performed in the R package ‘adespatial’ using the function *dbmem* which first calculates a Euclidean distance matrix among all populations. The threshold value *thresh* for the truncation of the geographic distance matrix was set as the length of the longest edge of the minimum spanning tree. All distances that were larger than the truncation threshold were modified to 4\**thresh*. A Principal Coordinates Analysis (PCoA) was then performed on the modified matrix and eigenfunctions that model positive spatial correlation (Moran’s *I* larger than expected value of *I*) of the dbMEMs were retained as spatial variables. A forward selection procedure was performed to reduce the number of dbMEMs (Blanchet, Legendre, & Borcard, 2008).

Environmental variables were derived from categorical (mountains and rivers; Fig. 2A, B) and continuous raster data (forest cover and habitat suitability; Fig. 2C, D). To test the Mountain Range Hypothesis, populations from the same mountain ranges were assumed to exchange genes freely and were given a low resistance value of 1. Populations from different but adjacent mountain ranges were given a value of 10 and populations from non-adjacent mountain ranges (e.g. between western and eastern ranges) were given a value of 100 (Fig. 2A). These values were selected because of the limited dispersal abilities of *Amolops* that have been shown to move freely within its own stream network, but does not disperse across intervening unforested lowlands that separate the mountain ranges (Chan *et al*., 2018a). Because Torrent Frogs are strictly associated with streams, an independent landscape variable based on river basin configurations was also constructed. River basins were calculated from a 90 m SRTM digital elevation model using the QGIS GRASS plugin *r.watershed* (QGIS Development Team 2017) with a minimum size of exterior watershed basin set at 10,000. Similarly, populations that occurred within the same major river basin were given a low resistance value of 1 and populations that occurred in separate basins were given a resistance value of 10 (Fig. 2B).

**Fig. 2.**
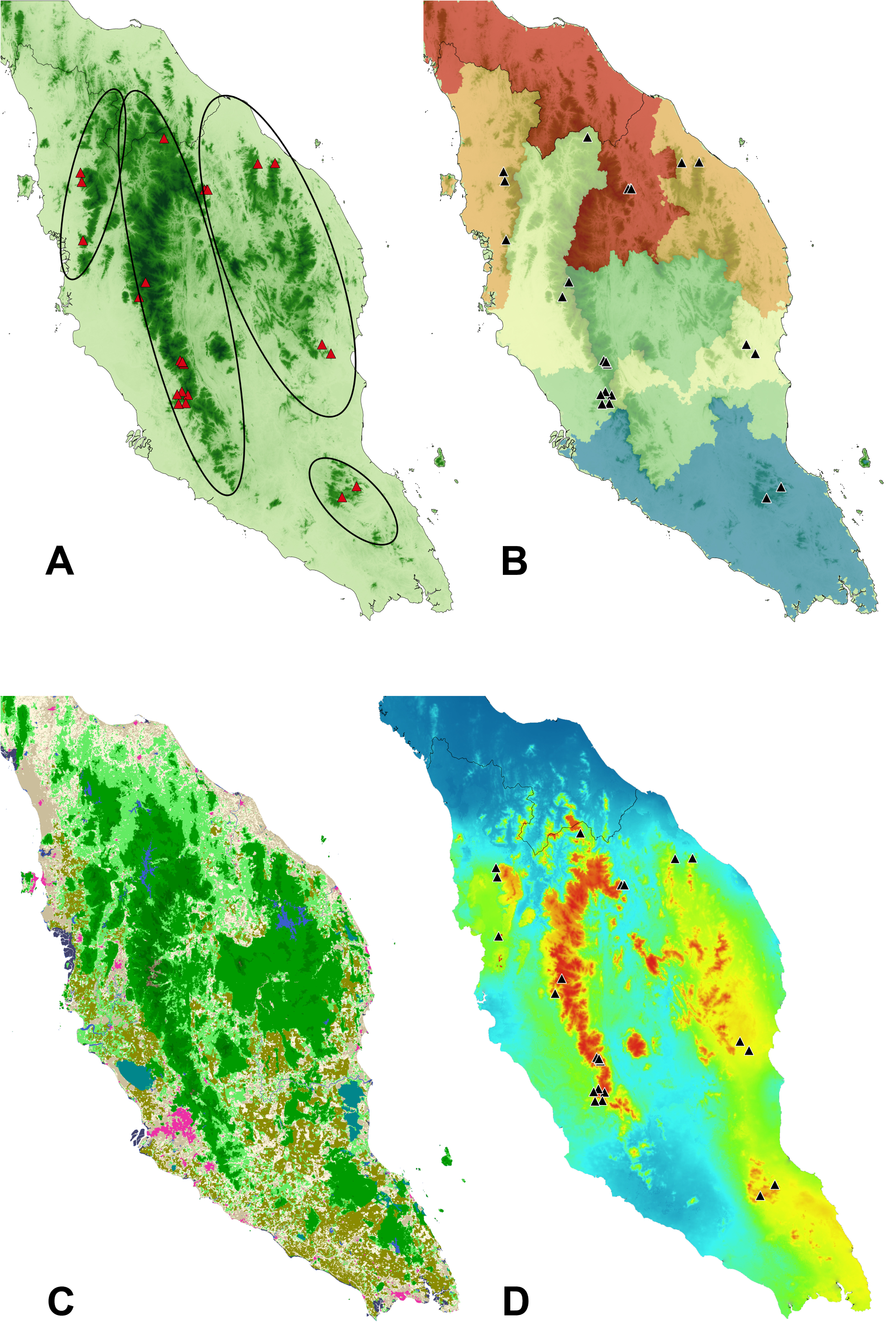
Environmental layers used to generate resistance matrices in Circuitscape: (A) Mountains; ellipses show the approximate boundaries of the four considered mountain ranges; (B) River basins; colors represent major watersheds estimated using QGIS; (C) Forest cover; raster layer of land use; (D) Habitat suitability; environmental niche model derived from the first three principal components of 19 Bioclim variables.

For continuous landscape variables, raster data for forest cover were derived from a 2015 land cover map of Southeast Asia at 250 m spatial resolution (Miettinen, Shi, & Liew, 2016) obtained from the Centre of Remote Imaging, Sensing, and Processing (CRISP) at the National University of Singapore (Fig. 2C). Habitat suitability was represented by an ecological niche model generated using 19 WorldClim (version 2) bioclimatic variables at a resolution of 30 seconds (Fick & Hijmans, 2017). To reduce dimensionality and correlation among variables, climatic variables were subjected to a Principal Components Analysis using the R function *iPCARaster* from the package ‘ENMGadgets’ (Barve & Barve, 2014). The first three principal components which accounted for 99% of the total variance were retained and used to generate niche models in the program Maxent (Phillips *et al*., 2006). The final niche model was constructed using the median values of 10 independent Maxent runs (Fig. 2D). All categorical and continuous landscape variables were subsequently transformed into resistance matrices using CIRCUITSCAPE 4.0 (McRae & Beier, 2007; McRae & Shah, 2009). To test for IBC, an east-west ancestry variable that corresponded to the major diverging split of the eastern and western lineages was used (Nadeau *et al*., 2016). This variable was represented by the Q-values from the sNMF population structure analysis at K=2 (Chan *et al*., 2017).

### Landscape genomic analyses

Mantel tests are one of the most widely-used approaches to assess spatial processes that drive population structure, particularly to detect IBD and to parse out the relative contributions of IBD and IBE (Meirmans, 2012). However, numerous studies have demonstrated that Mantel tests do not provide an accurate decomposition of genetic variation and therefore are unable to detect spatial structures or control for spatial autocorrelation in the relationship between genetic variation (Meirmans, 2012; Diniz-Filho *et al*., 2013; Guillot & Rousset, 2013; Legendre *et al*., 2015). As an alternative, ordination techniques such as redundancy analysis (RDA) was shown to be an improvement over the Mantel test because they do not require distance-based metrics and can overcome underlying assumptions of the Mantel test (e.g. linear relationships between variables). Additionally, distance-based RDA (dbRDA) can utilize transformations such as principal coordinates analysis (PCoA) to linearize genetic variables, thus removing any potential violations of linearity observed in Mantel tests (Guillot & Rousset, 2013; Kierepka & Latch, 2015). Furthermore, RDA is able to provide ANOVA-like statistics such as estimates of variance around F-ratios and contributions of each dependent variable, which allows for more robust interpretation of results compared to Mantel tests that only provide a correlation coefficient and P-value (Kierepka & Latch, 2015; Legendre *et al*., 2015). Given these advantages, constrained ordination methods such as dbRDAs represent more robust, and methodologically improved alternatives to formerly widely adopted spatial analyses of population structure.

A dbRDA analysis was implemented to assess the significance and relative contribution of IBD, IBE, and IBC to genetic variation. Because dbRDA requires independent variables to be site-specific, the pairwise resistance matrices from CIRCUITSCAPE were summarized by population using a negative exponential distribution kernel that represents a connectivity index based on the Incidence Function model:

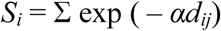

*S_i_* is the connectivity index for the cell *i*, *α* is a scalar correlated with average dispersal distance of the species, and *d* is the resistance value between sites *i* and *j* (Moilanen & Nieminen, 2002; Kierepka *et al*., 2016). Because no empirical data is available on the average dispersal distance of *Amolops*, *α* was set to a value of 1. We consider this value to be reasonable given the similarity in ecology/behavior of *Amolops* that does not suggest different average dispersal distances among populations or species. All independent variable matrices were scaled to a mean of zero and a variance of one prior to dbRDA analyses. The dependent variable (genetic distance) was left untransformed because the dbRDA analysis transforms this variable using PCoA and extracts all PCoA vectors that have positive eigenvalues. To inspect the potential correlation between independent variables, a correlation analysis was also performed. For brevity, the independent variables will be referred to as Distance (dbMEMs of geographic distance), Ancestry (east-west Q-matrix), and Mountains, Rivers, Forest, and Habitat for the resistance matrices representing mountain ranges, river basins, forest cover and habitat suitability respectively.

The dbRDA analysis was first performed on the entire dataset that included both the western and eastern lineages. A potential problem that could arise from analyzing datasets which include a large range of genetic divergences is that strong signals (e.g. diversification of the western and eastern lineages) could potentially overwhelm weaker signals which could be acting independently on populations within the constituent lineages. Additionally, assessing the contribution of specific variables responsible for within- or between-lineage diversification would be problematic due to non-identifiability issues inherent in a hierarchical dataset. To circumvent these problems, the dbRDA analysis was also performed on separate datasets consisting of eastern and western populations exclusively.

The dbRDA analysis was first performed separately on each independent variable using the R function *capscale* in the package ‘vegan’ (Oksanen *et al*., 2017). To parse out the effects of spatial autocorrelation, a partial ordination analysis was subsequently performed by conditioning each variable on geographic distance. Statistical significance of models was assessed using 999 permutations. In addition to the R^2^ statistic, adjusted R^2^ (R^2^_adj_) was also calculated to adjust for multiple predictors in the model. The best overall model was assessed using a forward and backward selection procedure (Blanchet *et al*., 2008) to select the most significant variables that explained the observed genetic variation. To avoid including correlated variables in the model, the variance inflation factor (VIF) statistic was calculated, where a VIF value of 1 represents completely independent variables, and values above 10 are regarded as highly multicollinear (Oksanen, 2012). Finally, we used variance partitioning (Borcard, Legendre, & Drapeau, 1992) from the R function *varpart* to provide a neutral decomposition of variation into unique and shared components using environmental, spatial and ancestral components as sources of genetic variation.

The concept of effective migration was used to model and visualize the relationship between genetics and geography using an estimated effective migration surface approach, EEMS (Petkova, Novembre, & Stephens, 2016). Briefly, a triangular grid was overlaid onto the study region and each sampling point was assigned to the closest deme on the grid. Using a stepping-stone model, each deme can only exchange migrants with its neighbors. Expected genetic dissimilarities depend on sample locations and migration rates and the expected genetic dissimilarity between two individuals were computed by integrating over all possible migration histories in their genetic ancestry and then approximated using resistance distance that integrates all possible migration routes between two demes. The estimation procedure adjusts migration rates for all edges so that genetic differences expected under the model matches the observed genetic differences from the data. These estimates were then interpolated across the habitat to produce an estimated effective migration surface, EEMS (Petkova *et al*., 2016).

For the EEMS analysis, the genetic dissimilarity matrix was computed using the mean allele frequency imputation method implemented in the R script ‘str2diffs’ available from the EEMS GitHub repository. We used two different grid densities (200 and 300 demes) to assess sensitivity to grid resolution on the interpolation of effective migration rates across a landscape. Four independent MCMC chains (500,000 generations/chain) were executed from different randomly initialized parameter states. Chains were combined and assessed for convergence by plotting the trace of log posterior probabilities against MCMC iterations after a 100,000 generation burn-in period.

### Population demographic analyses

Demographic history was estimated using a stairway plot approach which infers changes in population size over time using SNP frequency spectra, SFS (Liu & Fu, 2015). To reduce the effects of missing data and computational time, the SNP dataset was filtered to include only loci that contained no more than 10% missing data. As input into the stairway ploy analysis, a folded SFS was generated using δaδi v1.7.0 (Gutenkunst *et al*., 2009). The total number of observed nucleic sites (including polymorphic and monomorphic) was calculated from raw pyRAD outputs and set at 10,000,000, while mutation rate was set at 1.9 × 10^−8^ (Chan *et al*., 2017). We used 67% percent of sites for training with four random break points for each try (18, 36, 54, and 70). Stairway plot estimations were generated using 200 bootstrap SFSs.

To estimate lineage-specific changes in contemporary and ancestral population sizes, we implemented a Generalized Phylogenetic Coalescent Sampler (G-PhoCS) approach (Gronau *et al*., 2011). Demographic parameters were estimated for all three species and priors for all population size (θ) and divergence time (τ) parameters used a gamma distribution with α = 1, β = 10,000 and priors for all migration (*m*) parameters used a gamma distribution with α = 0.002 and β = 0.00001 following recommendations from (Gronau *et al*., 2011). A constant rate was used to model rate variation across loci and sampling was performed using a total of 300,000 MCMC iterations with the first 100,000 discarded as burn-in. The automatic fine-tune procedure was invoked and fine-tune settings were dynamically searched using the first 10,000 samples, with fine-tunes being updated every 100 samples. The program Tracer (Rambaut *et al*., 2014) was used to assess MCMC convergence and summarize parameter estimates.

## RESULTS

### IBD, IBE, and IBC

Results of the dbRDA analyses are summarized in Table 1: Distance and Ancestry are the only variables that show significant associations with genetic distance (*p* = 0.001 and 0.008 respectively). This is also reflected in high R^2^_adj_ values (R^2^_adj_ = 0.67 and 0.55 respectively). The significant correlation between Ancestry and genetic distance remains significant after separating out the effect of Distance [*p*(Ancestry | Distance) = 0.04]. None of the environmental variables show significant relationships with genetic distance with the exception of Rivers that is marginally significant [*p*(Rivers) = 0.05]. However, Rivers becomes insignificant after controlling for Distance [*p*(Rivers | Distance) = 0.4]. When the dbRDA analysis was performed separately on the western dataset, Distance remain significant (*p* = 0.001) while Ancestry is not (*p* = 0.08). There is no relationship between genetic distance and Ancestry after controlling for Distance [p(Ancestry_West_ | Distance = 0.9; R^2^_adj_ = −0.04). For the eastern dataset, Distance and Ancestry are significant (*p* = 0.008 and 0.05 respectively) and Ancestry remain significant after controlling for Distance [*p*(Ancestry_East_ | Distance = 0.02; R^2^_adj_ = 0.6). Both forward and backward selection procedures select Distance and Ancestry as the only variables that significantly explain genetic variation in the dataset, while Habitat contributes the least (Table 2).

**Table 1.**
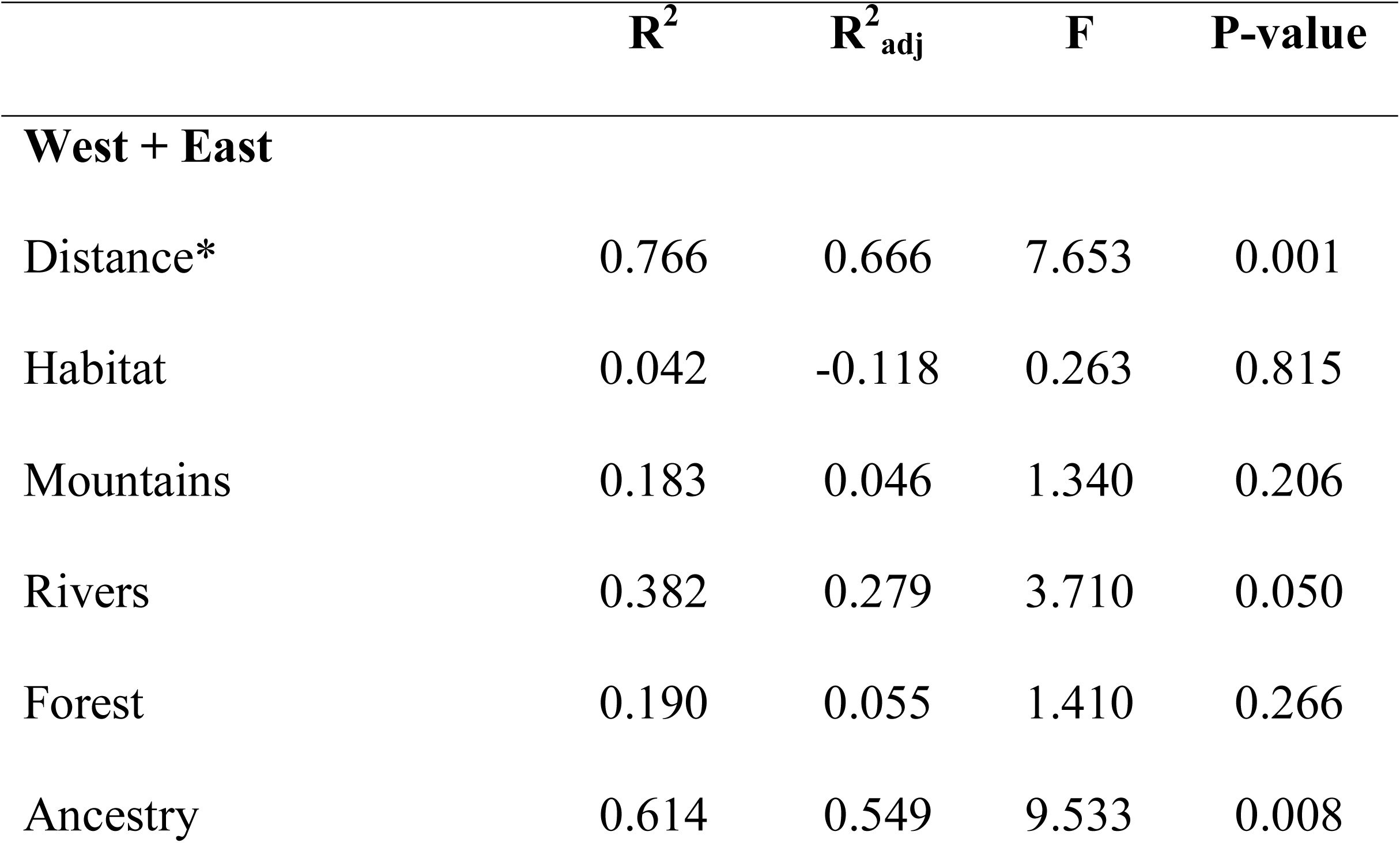
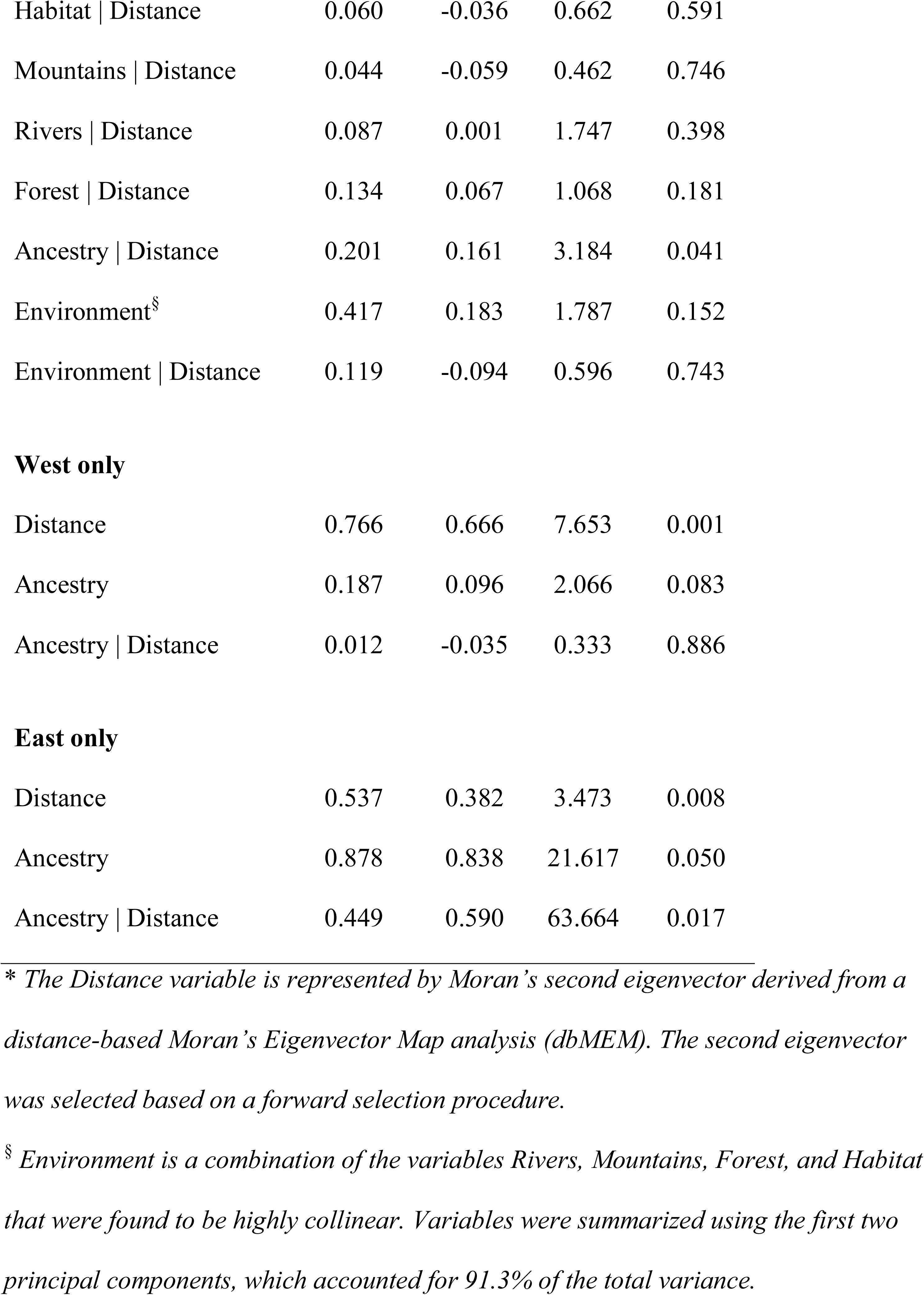
Results of the distance-based redundancy analysis (dbRDA). Effects of variables after “|” have been controlled for in the analysis.

| | $R^2$ | $R^2_{adj}$ | F | P-value |
| --- | --- | --- | --- | --- |
| **West + East** |  |  |  |  |
| Distance* | 0.766 | 0.666 | 7.653 | 0.001 |
| Habitat | 0.042 | -0.118 | 0.263 | 0.815 |
| Mountains | 0.183 | 0.046 | 1.340 | 0.206 |
| Rivers | 0.382 | 0.279 | 3.710 | 0.050 |
| Forest | 0.190 | 0.055 | 1.410 | 0.266 |
| Ancestry | 0.614 | 0.549 | 9.533 | 0.008 |
| Habitat \| Distance | 0.060 | -0.036 | 0.662 | 0.591 |
| Mountains \| Distance | 0.044 | -0.059 | 0.462 | 0.746 |
| Rivers \| Distance | 0.087 | 0.001 | 1.747 | 0.398 |
| Forest \| Distance | 0.134 | 0.067 | 1.068 | 0.181 |
| Ancestry \| Distance | 0.201 | 0.161 | 3.184 | 0.041 |
| Environment<sup>§</sup> | 0.417 | 0.183 | 1.787 | 0.152 |
| Environment \| Distance | 0.119 | -0.094 | 0.596 | 0.743 |
| **West only** |  |  |  |  |
| Distance | 0.766 | 0.666 | 7.653 | 0.001 |
| Ancestry | 0.187 | 0.096 | 2.066 | 0.083 |
| Ancestry \| Distance | 0.012 | -0.035 | 0.333 | 0.886 |
| **East only** |  |  |  |  |
| Distance | 0.537 | 0.382 | 3.473 | 0.008 |
| Ancestry | 0.878 | 0.838 | 21.617 | 0.050 |
| Ancestry \| Distance | 0.449 | 0.590 | 63.664 | 0.017 |

\* The Distance variable is represented by Moran's second eigenvector derived from a distance-based Moran's Eigenvector Map analysis (dbMEM). The second eigenvector was selected based on a forward selection procedure.
<sup>§</sup> Environment is a combination of the variables Rivers, Mountains, Forest, and Habitat that were found to be highly collinear. Variables were summarized using the first two principal components, which accounted for 91.3% of the total variance.

**Table 2.**
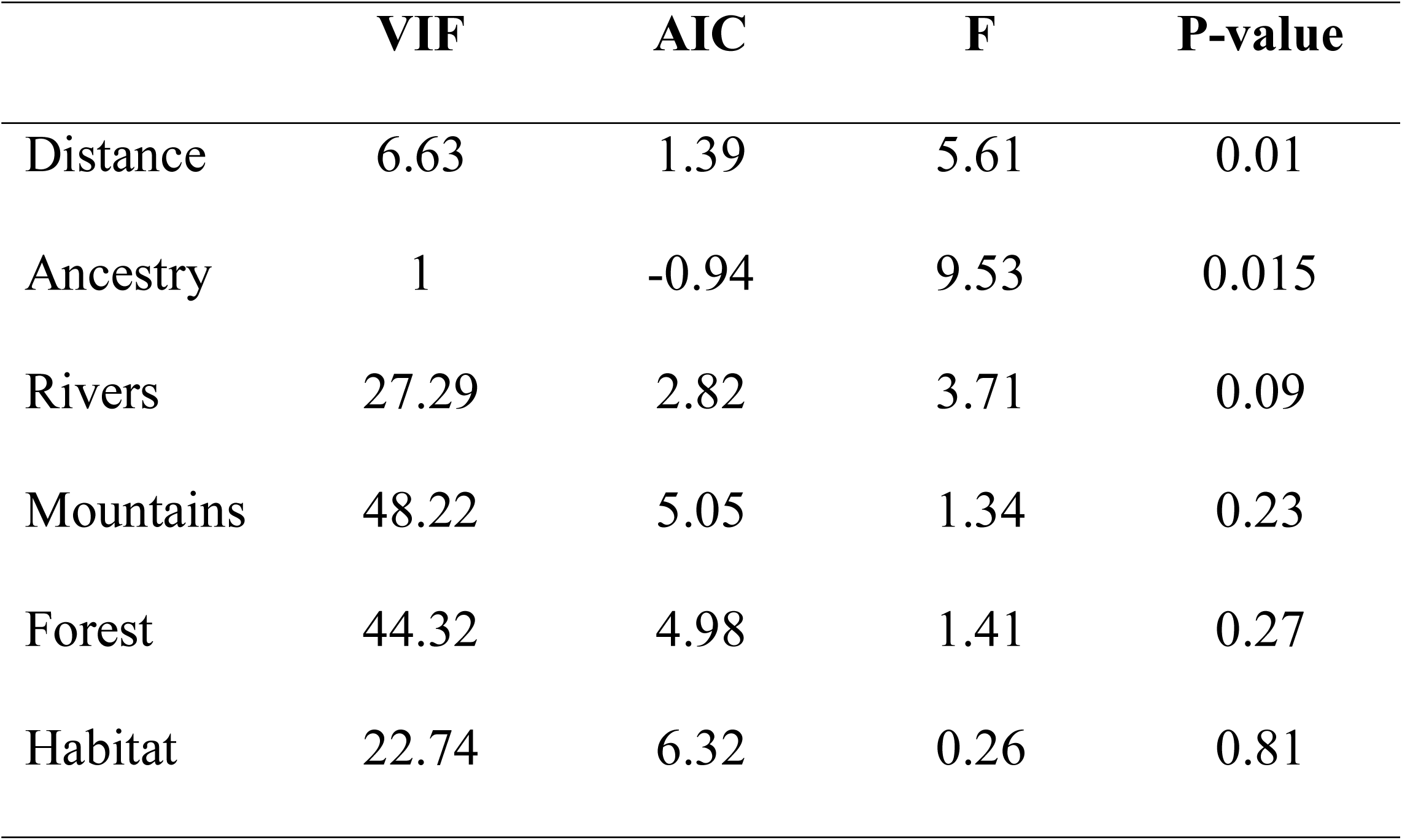
Results of model testing using the forward selection procedure. VIF = variance inflation factor; AIC = Akaike information criterion.

|  | VIF | AIC | F | P-value |
| --- | --- | --- | --- | --- |
| Distance | 6.63 | 1.39 | 5.61 | 0.01 |
| Ancestry | 1 | -0.94 | 9.53 | 0.015 |
| Rivers | 27.29 | 2.82 | 3.71 | 0.09 |
| Mountains | 48.22 | 5.05 | 1.34 | 0.23 |
| Forest | 44.32 | 4.98 | 1.41 | 0.27 |
| Habitat | 22.74 | 6.32 | 0.26 | 0.81 |

A Pearson’s correlation test (at α = 0.01) showed that Rivers and Mountains are significantly positively correlated while Ancestry and Distance are negatively correlated (Fig. 3). A further assessment of multicollinearity among independent variables within the RDA model testing framework was performed using the variance inflation factor (VIF). Results show that Ancestry and Distance are the only variables that are independent (VIF < 10) while the variables Rivers, Mountains, Forest, and Habitat are highly collinear (VIF > 20; Table 2). Because multicollinearity of independent variables increases estimates of parameter variance that can result in failure to detect significance in the model, these collinear variables were combined into a single variable, “Environment.” The dimensionality of the Environment variable was reduced using PCA and the first two PCs that accounted for 91.3% of the total variance were retained. The dbRDA analysis was then performed on the retained PCs but the results remain insignificant [*p*(Environment) = 0.15; R^2^_adj_ = 0.18] (Table 1).

**Fig. 3.**
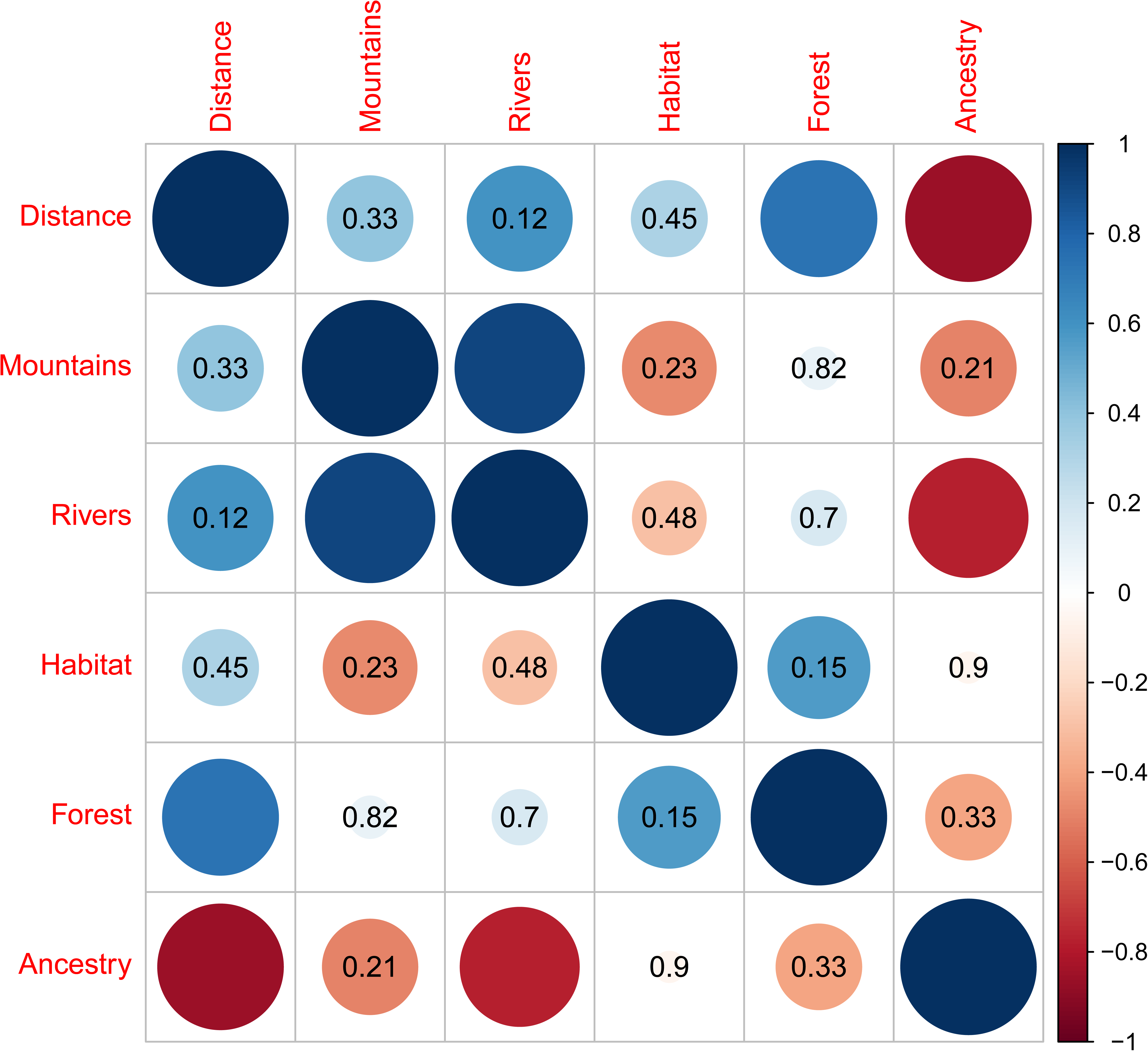
Pairwise correlation plot of independent variables and corresponding P-values. Circles without values represent *p* < 0.01.

Using R^2^_adj_ values from the RDA analyses, genetic differentiation was partitioned into three components: (1) Distance (IBD) represented by “Distance;” (2) Environment (IBE) represented by the combination of the Mountains, Rivers, Forest, and Habitat variables; and (3) Ancestry (IBC) represented by the east-west ancestry variable “Ancestry.” A total of 78% of the variation in Peninsular Malaysian *Amolops* can be explained by the three components and their various combinations. Ancestry contributes 23% after controlling for Environment and Distance, while Environment and Distance by themselves did not contribute to overall variation. A total of 70.5% of the explained variation is confounded between the effects of Distance, Environment and Ancestry (Fig. 4A). When the western and eastern lineages were analyzed separately, Distance and Ancestry explain 79% of the total variation within the western lineage. However, Distance contributes to most of this variation (53%) while Ancestry contributes 0%. A total of 16% is confounded by a shared effect between Distance and Ancestry (Fig. 4B). Conversely for the eastern lineage, Distance only explains 12% while Ancestry explains most of the variation (53%) (Fig. 4C).

**Fig. 4.**
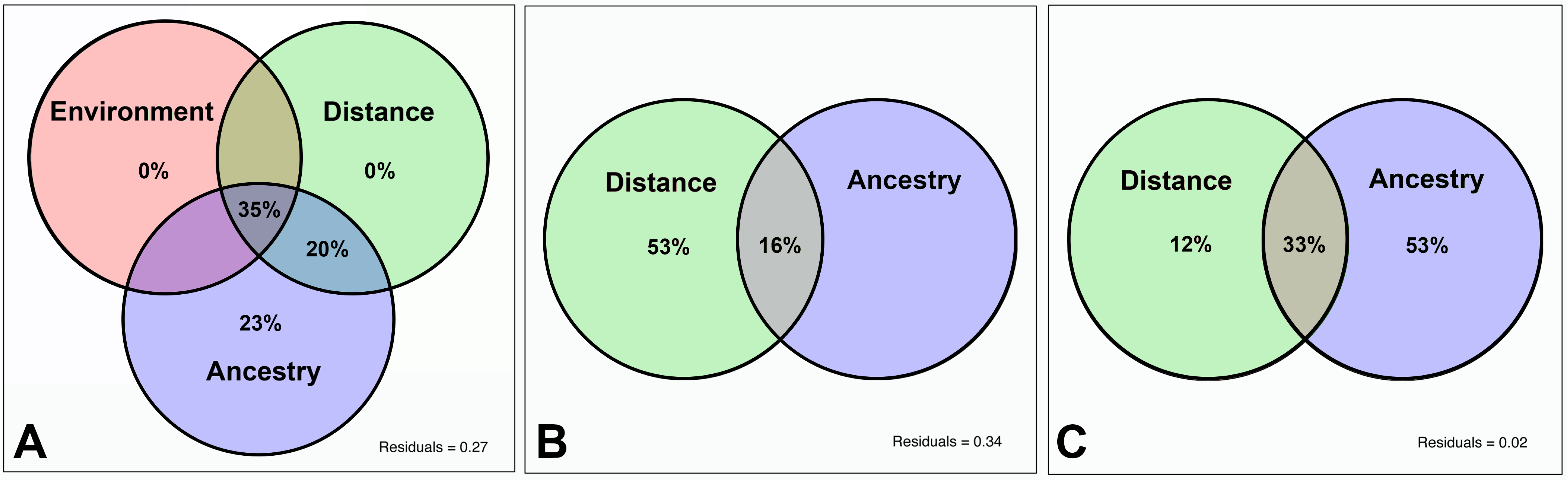
Venn diagrams showing the relative contributions of Environment, Distance and Ancestry to genetic distance. (A) whole dataset; (B) western lineage only; (C) eastern lineage only.

### EEMS and demographic history

All MCMC chains for the EEMS analysis achieved convergence (Fig. S1). Plots of observed versus fitted dissimilarities between and within demes show a strong linear relationship (Fig. S1), indicating that the EEMS model fits the data well. As expected, the analysis using 300 demes shows better resolution and detects finer-scale differences in migration and diversity estimates and is therefore retained for subsequent discussions. The estimated effective migration surface derived from posterior mean migration rates (*m*) shows that effective migration rates are higher than the overall average (highlighted in blue) among populations within the western (*Amolops larutensis*) and eastern lineages (*A. gerutu* + *A. australis*). Conversely, effective migration rates are lower than the overall average (highlighted in orange) in areas separating eastern from western populations and also separates the southern population from both eastern and western populations (Fig. 5A). Areas where effective migration rates are significantly higher or lower than the overall average can be emphasized by highlighting areas where the posterior probability Pr{*m* > 0} or Pr{m < 0} exceeds 90% and can be interpreted as areas representing corridors and barriers to gene flow (Fig. 5B). Diversity estimates are congruent with migration rate estimates where corridor areas along the northwestern mountain range show low diversity estimates (orange), while areas along the eastern and southern mountain ranges show higher diversity estimates (blue) (Fig. 5C).

**Fig. 5.**
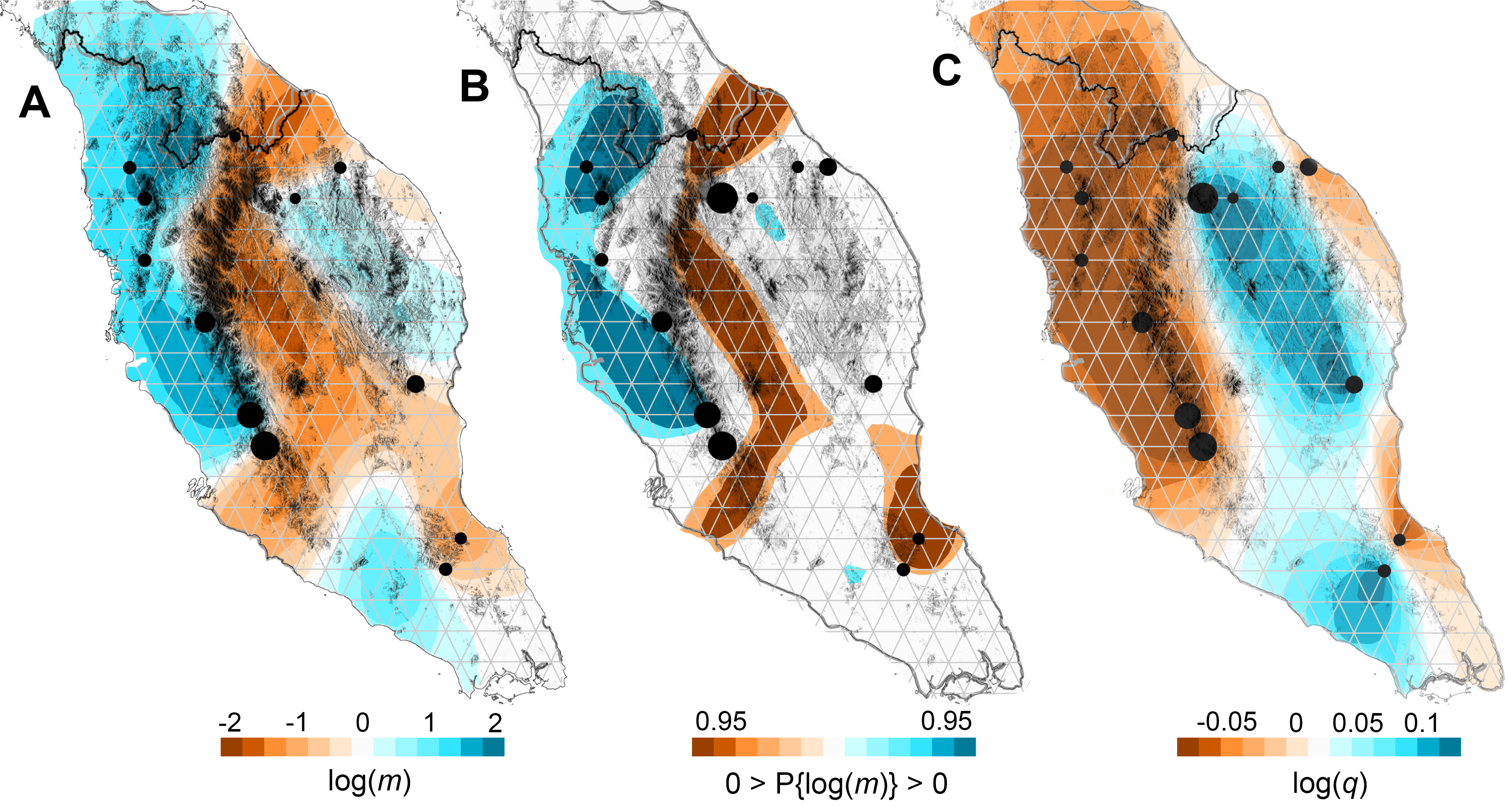
Estimated effective migration surfaces (EEMS) at 300 demes. (A) Posterior mean migration rates (*m*) on a log_10_ scale after mean centering, where 0 corresponds to the overall mean migration rate; (B) Plot emphasizing regions where effective migration rates are significantly higher/lower than overall average. Areas where the posterior probability Pr{*m* > 0} exceeds 90% are highlighted in blue, areas where Pr{*m* < 0} exceeds 90% highlighted in orange; (C) Posterior mean diversity rates (*q*) on a log_10_ scale after mean centering.

The stairway plot analysis reveal two bottleneck events that severely reduces the overall effective population size of *Amolops* (Fig. 6). These results are also supported by the G-PhoCS analysis. The ancestor of Peninsular Malaysia *Amolops* has a large population size (θ_root_ = 120) that was drastically reduced when the western lineage (*A. larutensis*) diverged (θ_west_ = 44). Similarly, the ancestral population size of the eastern lineage (θ_ES_= 107) was severely reduced when *A. gerutu* and *A. australis* diverged (θ_east_ = 17; θ_South_ = 5) (Table 3; Fig. 6). Most parameters converge with high ESS values (>500) with the exception of θ_west_ and θ_east_ that had moderate ESS values (200 > ESS > 150; Table 3). These bottleneck events are corroborated by low observed heterozygosity measures for the tree major lineages: west, east, and south (Fig. 6; Table 3).

**Fig. 6.**
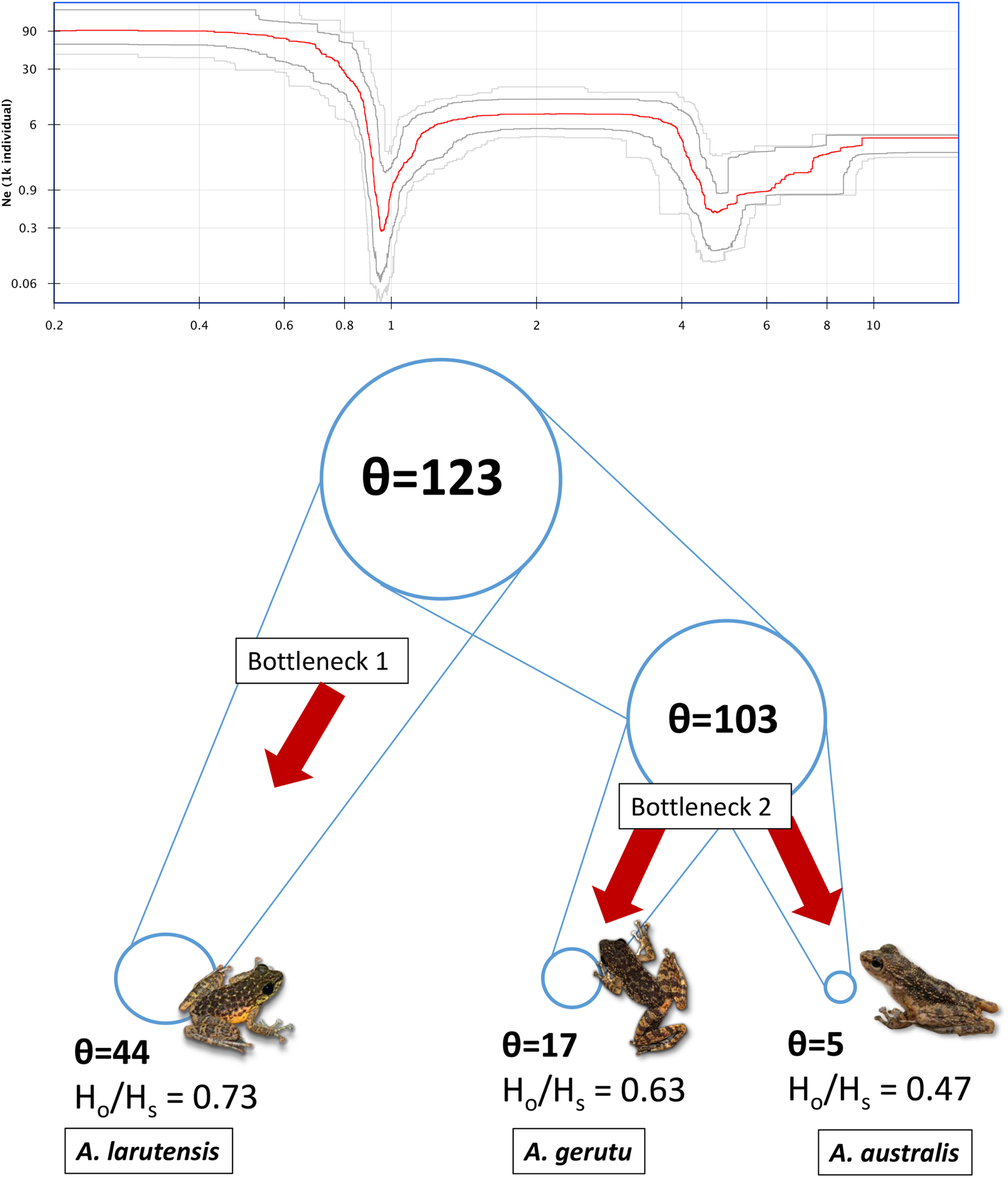
Stairway plot showing changes in effective population size (N_e_) through relative time (top); bottlenecks and heterozygosity measures for the three species of Malaysian torrent frogs (bottom).

**Table 3.** **Top:** Genetic diversity estimates for the three species of *Amolops* in Peninsular Malaysia. Num = number of alleles; Eff. Num = effective number of alleles; H_o_ = observed heterozygosity; H_s_ = expected heterozygosity; G_is_ = inbreeding coefficient. **Bottom:** G-PhoCS summary statistics of population size parameter (θ) estimated for contemporary western (θ_west_), eastern (θ_east_), southern (θ_south_); and the ancestor of the east + south populations (θ_ES_) and ancestor of the entire clade (θ_root_). Parameter estimates are relative values that not scaled to represent effective or consensus population sizes. SE = standard error, Std Dev = standard deviation.

| Gen. Diversity | Num | Eff. Num | $H_o$ | $H_s$ | $G_{is}$ |
| --- | --- | --- | --- | --- | --- |
| *A. larutensis* | 1.398 | 1.027 | 0.016 | 0.022 | 0.262 |
| *A. gerutu* | 1.269 | 1.044 | 0.020 | 0.032 | 0.384 |
| *A. australis* | 1.153 | 1.080 | 0.027 | 0.058 | 0.533 |

| G-PhoCS | $\theta_{\text{west}}$ | $\theta_{\text{east}}$ | $\theta_{\text{south}}$ | $\theta_{\text{ES}}$ | $\theta_{\text{root}}$ |
| --- | --- | --- | --- | --- | --- |
| Mean | 43.74 | 16.61 | 4.73 | 102.60 | 122.64 |
| SE mean | 0.022 | 0.0456 | 0.0156 | 0.0512 | 0.1251 |
| Std Dev | 0.5516 | 0.5466 | 0.1926 | 2.1486 | 4.3437 |
| Variance | 0.304 | 0.2988 | 0.0371 | 4.6165 | 18.8681 |
| Median | 43.734 | 16.60 | 4.73 | 102.58 | 122.64 |
| Geometric mean | 43.73 | 16.60 | 4.73 | 102.58 | 122.57 |
| 95% HPD | [42.65, 44.81] | [15.55, 17.65] | [4.36, 5.11] | [98.47, 106.89] | [114.13, 131.08] |
| ESS | 631.32 | 151.54 | 191.93 | 1760.21 | 1205.30 |

## DISCUSSION

### Biogeography

Our findings did not support the hypothesis that mountain ranges have played a strong and direct role in the initial diversification of Malaysian *Amolops*. In fact, the distribution of river basins explained significantly more variation. This was not unexpected given the ecology of *Amolops* that is strictly associated with river systems. However, Mountains and Rivers were also shown to be highly correlated, most likely because calculations of river basins were predicated on a digital elevation model. These results show that mountain ranges per se should not be considered strictly as a physical barrier to gene flow, but instead be viewed as part of an interactive complex of environmental variables (including river basins) that can have confounding effects on genetic variation. It should also be noted that the values used to model dispersal resistance among mountains and river basins were arbitrary due to the lack of empirical data on the dispersal attributes of these species. The rejection of the mountain range hypothesis should therefore be interpreted with caution pending more robust analyses.

Traditional expectations view the western, central, and eastern mountain ranges (the southern range is usually considered as part of the eastern range) as separate biogeographic units (Fig. 2A) that harbor unique species assemblages (Grismer *et al*., 2012, 2013). Although this is likely true for some species, our results show that genetic structuring among populations from the western and central mountain ranges are a result of isolation-by-distance as opposed to isolation by mountain range. These results are in line with findings from a previous study that demonstrated gene exchange between western and central populations of *Amolops* (Chan *et al*., 2017), indicating the presence of habitat corridors between these ranges. However, our results show the presence of a strong genetic barrier between the central and eastern ranges, and between the central and southern ranges (Fig. 2A). These patterns of genetic isolation are also supported by numerous other studies (Grismer, 2007; Grismer *et al*., 2008, 2014a; Matsui *et al*., 2012; Matsui, Belabut, & Ahmad, 2014; Chan *et al*., 2014).

### Drivers of genetic differentiation

When western and eastern lineages were jointly analyzed, <u>Distance</u> significantly contributed to the overall genetic variation (Table 1). However, when the effects of Environment and Ancestry were controlled for, Distance by itself did not uniquely contribute to genetic variation but produced shared confounding effects with the other variables (Fig. 4A). Further insights into these confounding effects were gained when western (*Amolops larutensis*) and eastern lineages (*A. gerutu* + *A. australis*) were analyzed separately: for the western lineage, Distance was the main source of genetic variation (53%) whereas for the eastern lineage, Distance only contributed 12%, indicating that Distance had contrasting effects on the genetic variation of the two major lineages. The differential effects of Distance can be due to the physiographic differences of the different mountain ranges. Populations of *A. larutensis* mainly occurred along the contiguous central range that would have provided a continuous corridor for dispersal compared to the highly fragmented eastern range. Based on the current data, IBD explains most of the genetic variation among populations of *A. larutensis*; has a significantly smaller effect on populations of *A. gerutu* and *A. australis*; and produces confounding effects when both lineages are jointly analyzed. This highlights the importance of parsing-out analyses to subsets of data that could potentially yield misleading results when analyzed as a whole.

When environmental variables were analyzed separately, Rivers contributed the most, while Habitat played no role in shaping genetic diversity. When the effects of Ancestry and Distance were controlled for, none of the environmental variables, either considered separately or combined, contributed to the overall genetic variance, indicating that the environment played no role in shaping the genetic variation in *Amolops*. These results agree with the conservative ecology of *Amolops* that shows little to no environmental adaptation across species.

Ancestry was a significant component before and after accounting for spatial autocorrelation and contributed 23% of the genetic variation when western and eastern lineages were jointly analyzed (Fig. 4A). However, when the dataset was analyzed separately (western vs eastern), Ancestry was not significant universally, and did not contribute towards the genetic variation of western populations. In contrast, Ancestry was the main source of variation (53%) within the eastern lineage. These results suggest that IBC was responsible for the initial diversification event that separated *A. larutensis* and the MRCA of *A. gerutu* + *A. australis*, and subsequently for the diversification of *A. gerutu* and *A. australis*. This was followed by secondary contact between eastern and western populations at the contact zone (Fig. 1), which could explain the significant negative correlation between Ancestry and Distance (Fig. 3).

### Modes of speciation

Strong signatures of IBC associated with the diversification of *Amolops larutensis, A. gerutu*, and *A. australis*, coupled with insignificant contributions from IBE and IBD, indicate that these species could have diverged via vicariant or founder effect speciation (Mayr, 1942; Yeung *et al*., 2011; Runemark *et al*., 2012; Lawson *et al*., 2015). One of the theoretical expectations of a founder event is that effective population size changes through time (Orsini *et al*., 2013b). In a classic scenario, a small number of individuals are isolated from the source population, resulting in a bottleneck that accelerates the formation of novel allelic combinations via genetic drift (Mayr, 1954). Reduction in population sizes can also be caused by vicariance via range fragmentation (Perez *et al*., 2016). The stairway plot and G-PhoCS analyses detected two bottleneck events consistent with these scenarios and this was further corroborated by lower than expected heterozygosity measures and severe reductions in population sizes in all three species. However, these modes of speciation can be difficult to distinguish from one another because both processes involve an ancestral distribution which is split into discontinuous populations and subsequently prevented from exchanging genes (Mayr, 1963).

Unfortunately, time estimates from demographic analyses only represent relative values and we were unable to associate directly, the timing of the bottlenecks with the timing of diversification of these lineages. Absolute estimates of timing, and changes in population size, would require prior information on mutation rates and generation times, both of which were not available. However, the strong signatures of IBC, non-significant contributions of IBD and IBE, and population bottlenecks associated with species diversification, strongly suggests that interspecific diversification was a result of vicariance or founder effect speciation; and that intraspecific population structure was caused by IBD. The negligible effect of environmental factors indicates that natural selection may not play a significant role in the evolution of Malaysian *Amolops* species diversity. Assessing the role that founder events, selection, and/or environmental variation plays in speciation in other, co-distributed endemic riparian vertebrates of the Peninsular Biodiversity Hotspot remains an intriguing and important challenge for future studies.

## ACKNOWLEDGEMENTS

Field work was supported by a grant from the National Geographic Society to CKO and L. L. Grismer (Grant number 9722-15), multiple grants from the University of Kansas (KU) Biodiversity Institute Panorama Grant Program (to CKO), and genetic data were collected with support from NSF DEB 1702036 (to CKO) and a KU College of Liberal Arts and Sciences, Office of the Provost, Research Investment Council (RIC) Level II grant (Award No. 2300207) to RMB. We thank Paul Hime for providing critical comments prior to manuscript submission.

---

**Fig. S1.** Left: traceplots of the four independent MCMC chains from the EEMS analysis at 300 demes; Centre: fitted vs. observed dissimilarities between pairs of sampled demes; Right: fitted vs. observed dissimilarities within sampled demes.

