## Supplementary figures and images for "Linking patterns of genetic variation to processes of diversification in Malaysian torrent frogs (Anura: Ranidae: *Amolops*): a landscape genomics approach"

### Supplemental Figure S1

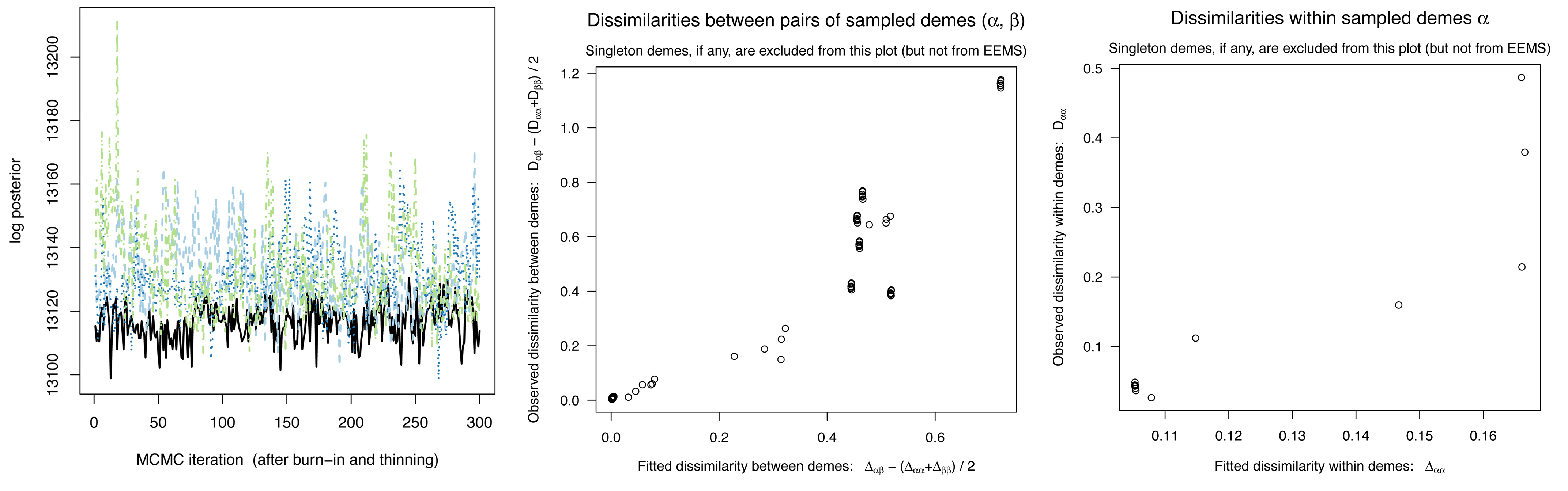
